# Learning a new class of multisensory associations: High-density electrophysiological mapping of the temporal course of audio-visual object processing

**DOI:** 10.1101/2021.11.15.468657

**Authors:** Tiziana Vercillo, Edward G. Freedman, Joshua B. Ewen, Sophie Molholm, John J. Foxe

## Abstract

Multisensory objects that are frequently encountered in the natural environment lead to strong associations across a distributed sensory cortical network, with the end result experience of a unitary percept. Remarkably little is known, however, about the cortical processes sub-serving multisensory object formation and recognition. To advance our understanding in this important domain, the present study investigated the brain processes involved in learning and identification of novel visual-auditory objects. Specifically, we introduce and test a rudimentary three-stage model of multisensory object-formation and processing. Thirty adults were remotely trained for a week to recognize a novel class of multisensory objects (3D shapes paired to complex sounds), and high-density event related potentials (ERPs) were recorded to the corresponding unisensory (shapes or sounds only) and multisensory (shapes and sounds) stimuli, before and after intensive training. We identified three major stages of multisensory processing: 1) an early, multisensory, automatic effect (<100 ms) in occipital areas, related to the detection of simultaneous audiovisual signals and not related to multisensory learning 2) an intermediate object-processing stage (100-200 ms) in occipital and parietal areas, sensitive to the learned multisensory associations and 3) a late multisensory processing stage (>250 ms) that appears to be involved in both object recognition and possibly memory consolidation. Results from this study provide support for multiple stages of multisensory object learning and recognition that are subserved by an extended network of cortical areas.

## INTRODUCTION

Neural systems have evolved an exquisite set of sensory organs/receptors to take advantage of the multiple forms of energy afforded by environmental objects and events (e.g. light, sound, smell, etc.), and these cues arriving at or actively sampled through the relevant sensory systems are integrated to create the most accurate and informative representation of the physical world (Stein, 1998; M. T. Wallace, Carriere, Perrault, Vaughan, & Stein, 2006). Initial models of when and where the brain performs this multisensory integration (MSI) were generally predicated upon a serial hierarchical framework, with processing assumed to occur first in dedicated unisensory cortical areas, followed by multisensory interactions and cross-sensory binding in higher-order association regions (Felleman & Van Essen, 1991). However, results from a multitude of neurophysiological studies have since provided countervailing evidence, demonstrating that MSI can occur even within primary sensory areas (Driver & Noesselt, 2008; John J. Foxe & Schroeder, 2005; Schroeder et al., 2003), and that these MSI processes can be recorded during the very earliest temporal stages of cortical processing. A temporal cascade of MSI effects then continues to affect processing through subsequent perceptual and cognitive timeframes (John J. Foxe & Schroeder, 2005; Molholm et al., 2002), involving a distributed network of brain regions (Cappe, Rouiller, & Barone, 2009; Davies-Thompson et al., 2018; Meijer, Mertens, Pennartz, Olcese, & Lansink, 2019). Still, the functional significance of these successive temporal frames of multisensory interactions across disparate cortical areas is not well understood. To date very little effort has been dedicated to examining the spatiotemporal cortical processes underlying how representations of multisensory objects are initially formed, and the influence of learning on these representations over time.

The brain regions and temporal dynamics of multisensory processing are influenced by the sensory modalities of the stimulus inputs, their functional significance (e.g., speech; tools), and the goals of the individual (Beauchamp, 2005; Lucan, Foxe, Gomez-Ramirez, Sathian, & Molholm, 2010; Macaluso, George, Dolan, Spence, & Driver, 2004; Talsma, Senkowski, Soto-Faraco, & Woldorff, 2010). Electroencephalographic (EEG) studies in both animals and humans have shown initial cortical multisensory interactions in early sensory regions, that coincide with the onset of cortical sensory processing (~50 ms poststimulus onset; John J. Foxe et al., 2000; Kayser, Petkov, & Logothetis, 2008; Molholm et al., 2002; Schroeder et al., 2003). These early interactions most likely reflect the initial tagging of multiple sensory inputs that potentially belong to the same object, as well as the multisensory driven enhancement of sensory representations (Fiebelkorn, Foxe, & Molholm, 2010; Mercier & Cappe, 2019).

Subsequent multisensory interactions emerge within higher order sensory areas that are typically engaged in more complex cortical functions, such as object representation. The lateral occipital complex (LOC) of the visual ventral stream is comprised of a system of higher-order sensory-perceptual processing regions that are strongly linked to visual object processing (Goodale et al., 1994; Goodale & Milner, 1992; Kalanit Grill-Spector, 2003; Sehatpour *et al*., 2006), and represents a strong candidate region for multisensory object processing and representation, as suggested by several neuroimaging studies (A. Amedi, Von Kriegstein, Van Atteveldt, Beauchamp, & Naumer, 2005; Erdogan, Chen, Garcea, Mahon, & Jacobs, 2016; Lucan et al., 2010). In support of this idea, electrophysiological studies in humans described a modulation of the visual N1 ERP component during audiovisual object presentation (Giard & Peronnet, 1999; Molholm et al., 2002). This is a negative-going visual component occurring ~150 ms after stimulus presentation that modulates as a function of object type (e.g. faces versus houses). The N1 has been typically localized within ventral visual areas (Shpaner, Molholm, Forde, & Foxe, 2013; Vogel & Luck, 2000), and reflects processing in higher order visual areas associated with specific object types (e.g., the fusiform face area; the parahippocampal place area; the fusiform body area (Kanwisher, McDermott, & Chun, 1997; Murray et al., 2002; Peelen & Downing, 2005; Rossion, Kung, & Tarr, 2004). As such, a subsequent stage of MSI wherein multisensory object information is combined to influence object-level representations may occur in the timeframe of the visual N1, around 150 ms post stimulus onset, and within the ventral visual pathway.

Very little is yet known about how these multisensory object representations are formed, and it is the characterization of this stage of multisensory processing that we seek to contribute to with the current study. We suggest that when sensory elements have no known relationship to each other, they would not evoke integrative multisensory responses during the perceptual phase of processing related to object-recognition, but rather, novel instances of multisensory inputs will need to be further analyzed and the novel association consolidated. Only after repeated exposures and active recognition of a relationship between the inputs, strong and consolidated multisensory representations, associated with the meaning of the objects, can be encoded into cortex. This stage of memory consolidation likely engages a broad neural network composed of multiple brain regions (Alain, Woods, & Knight, 1998; Courtney, Ungerleider, Keil, & Haxby, 1997; Curtis & D’Esposito, 2003) from the hippocampus to sensory and associative areas (Sehatpour et al., 2008).

To advance understanding of how multisensory associations are represented, we have developed a rudimentary three-stage model (see figure 1a) of multisensory object-formation that makes clear predictions about the temporal windows in which we might expect to see multisensory effects. The model describes an early stage of integration, occurring between 40-90 ms after the onset of a multisensory input, which serves as a temporal coincidence detector and is necessary for multisensory binding. We expect this early stage of integration to be automatic and to engage early sensory cortices. A second temporal stage of integration specific to the processing of well-learned objects, is triggered by activity in sensory-perceptual cortical areas involved in object representation, and should emerge in a later temporal window between 120 and 200 ms post-stimulus onset. The third temporal multisensory stage (> 220 ms) involves a distributed network of cortical areas, such as the hippocampus, anterior temporal lobe and frontal regions, and in this model is only engaged when new multisensory associations are in the process of being established. Based on previous findings, our model predicts strong multisensory interactions during the first stage of processing for both novel and well-known multisensory stimuli. We expect multisensory interactions during the second stage of processing only in response to well-established cross-sensory as sociations (i.e. known multisensory objects), but not for entirely novel multisensory associations. We would expect temporal stage-3 processing to be present as new multisensory objects are learned, and to become unnecessary and attenuate as multisensory object representations are solidified in object processing regions of higher order sensory cortices, which we propose is represented by temporal stage-2 in our model.

**Figure 1:**
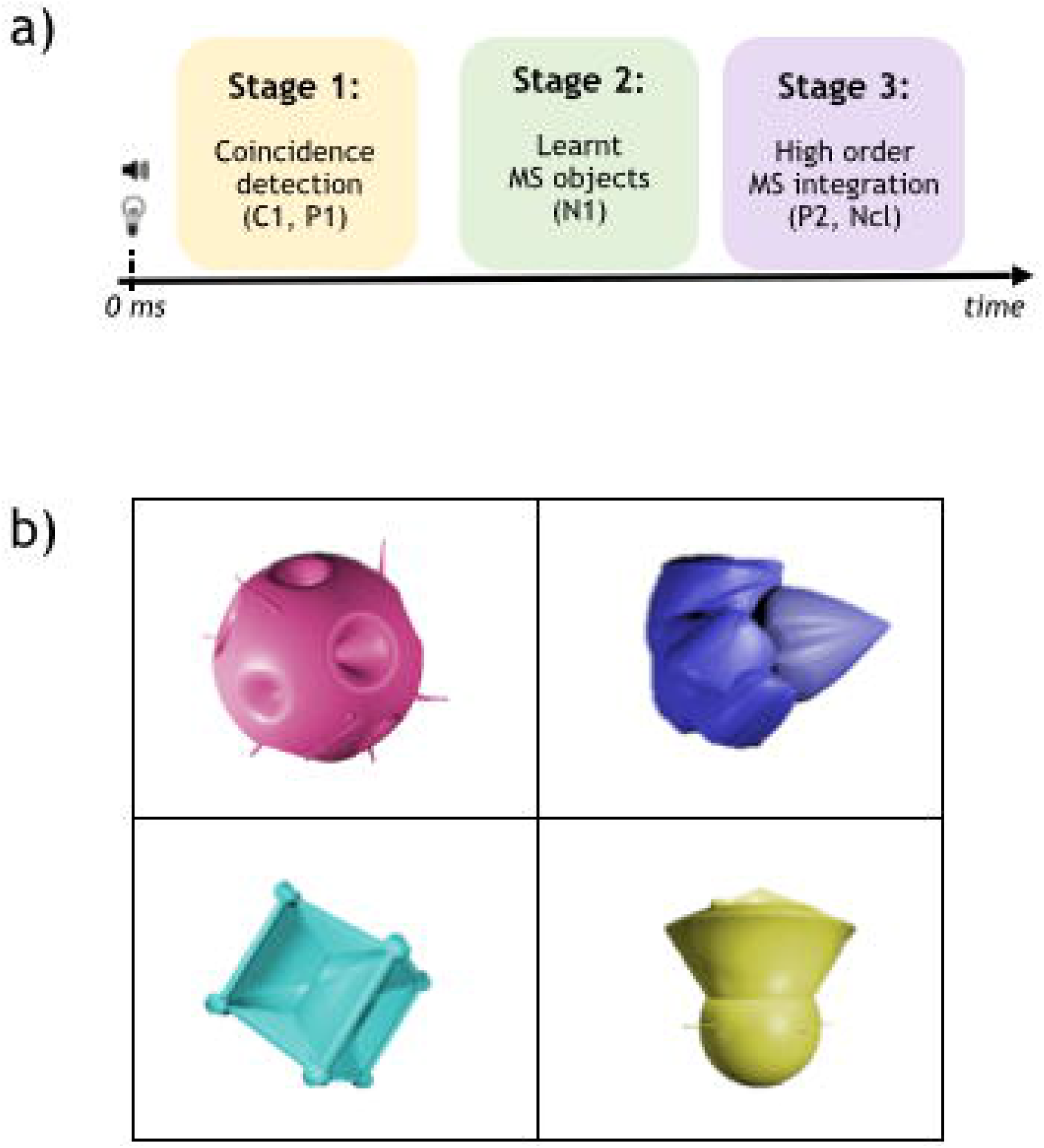
a) A schematic description of the model. The yellow box represents the first stage of coincidence detection occurring within 100 ms post-stimulus onset. The green box represents the second stage that is object specific and occurs around 100-150 ms post-stimulus onset. The purple box corresponds to the third stage and refers to the perceptual closure component that is visible from 200 ms after stimulus presentation b) Examples of the visual stimuli adopted in the study.

In the current study, we assess our 3-stage model and investigate the functional role of each successive temporal stage. Specifically, we examine whether MSI electrophysiological responses change over time in terms of their amplitude and the underlying brain network topologies, as the multisensory elements of a newly introduced class of audio-visual associations become more strongly bound into well-known object associations as a function of repeated exposures during an incentivized multi-day training protocol.

## METHODS

### Participants

Experimental procedures were approved by the Research Subjects Review Board at the University of Rochester and followed the principles of the Declaration of Helsinki. 30 young adults (23 ± 1 years old) participated in the study. All participants read and signed a written informed consent describing the experimental protocol, and received a modest payment for their participation.

### Visual, auditory and multisensory stimuli

The visual stimuli (see figure 1b for examples) were 2-D representations of twelve 3-D novel objects specifically designed to minimize subconscious associations with objects that exist. Each visual object was characterized by shape and color, with 12 possible colors selected from the color space and were separated by 30° (for all the visual shapes and colors see Supplemental Material, Supplemental Figure S1 - https://figshare.com/articles/journal_contribution/SupplementalMaterial_NOPA_Foxe_doc/19172378). The size of the visual stimuli was approximately 15X15 cm. Similarly, auditory stimuli were designed to avoid pre-existing associations. They consisted of 12 digitally fabricated sonic textures of 300 ms duration, within a comfortable frequency and volume range (70dB) presented from headphones (the spectrograms of all the auditory stimuli are displayed in Supplemental Figure S2). Multisensory objects were generated by pairing visual shapes with sounds, whereas colors varied randomly among the 12 possible hues. Stimulus delivery and timing was controlled through the stimulus delivery experiment control program “Presentation” (Neurobehavioral Systems, Inc).

### Experimental procedures

The experiment had three phases: pre-training EEG recording, intensive behavioral training (remote), and post-training EEG recording. The two EEG recordings shared the same paradigm and procedure.

During the EEG recordings, participants sat in a RF-shielded, sound attenuating booth at 57 cm from a computer screen (Acer, Predator Z35). Visual stimuli were displayed centrally on a grey background, while participants fixed their eyes on a yellow fixation cross presented at the center of the screen. Participants performed 17 blocks, each composed of 60 trials, for a total of 1020 trials (204 trials for experimental condition). Each block contained 12 unisensory visual trials, 12 unisensory auditory trials, 24 multisensory trials and 12 target trials. Unisensory visual/auditory trials consisted of the presentation of a visual/auditory stimulus alone for 300 ms duration. Multisensory trials involved the simultaneous presentation of a visual and an auditory stimulus for 300 ms duration. Trials from this condition were split into 12 “matches” and 12 “mismatches” (the distinction between these two categories is provided below, in the description of the training). Target trials were similar to multisensory trials, but the auditory or the visual stimulus was replaced by auditory (6 trials) or visual (6 trials) noise. The behavioral task was designed to maintain focused attention on both the auditory and the visual modalities throughout the duration of the EEG recording. Participants were asked to detect target trials (presence of auditory or visual noise) and to respond with a button press using their right index finger. Inter-stimulus-interval (ISI) was jittered between 800 to 1500 ms to avoid time-locked anticipatory responses, for methodological reasons (for rationale see e.g., Molholm et al., 2002). Stimulus sequence and randomization within each block was controlled through Matlab 2014a (The Mathworks, Inc.). The duration of each block was □ 2 min, and overall the duration of each EEG session was □ 40 min (participants were allowed to take breaks after each block). See Supplemental Figure S3 in the Supplemental Material for the EEG protocol - https://figshare.com/articles/journal_contribution/SupplementalMaterial_NOPA_Foxe_doc/19172378

During the training phase, participants engaged in a learning task entailing the memorization of 12 distinct shape-sound associations. After the first EEG session, participants received a tablet (Samsung Galaxy Tab E) to take home on which they were asked to play an interactive learning game on a daily basis, for one week. Training occurred over 21 sessions. Each session was comprised of a subject-directed exploration of the stimulus pairings and a test. Each multisensory object was presented for 300 ms and the color of the object was always randomized (see Supplemental Material for a detailed description of the training procedure - https://figshare.com/articles/journal_contribution/SupplementalMaterial_NOPA_Foxe_doc/19172378). Participants were free to explore the 12 objects at their own pace, for a total of 36 presentations (on average 3 times per object). After the exploration, participants’ learning was assessed in a 2 alternative forced choice (2AFC) task that was automatically delivered within the app. Two audiovisual objects were presented and participants were asked to identify the correct audiovisual pairing that matched one of the training audiovisual objects, i.e. the match. Overall, the task consisted of 12 trials. At the end of each training session performance feedback was provided. To motivate learning, participants were informed that they would receive a modest bonus for outstanding performance (10$-20$ for 80%-100% of correct responses for at least three consecutive sessions). Audiovisual pairings varied across participants randomly. Participants were required to perform 3 training sessions a day (with no temporal restriction) for 7 consecutive days. The app tracked the audiovisual objects considered as matches and the test results for each participant. This information was wirelessly transferred to a server for data analysis.

Phase 3, in which post-training EEG was recorded, occurred within 2 days from the completion of training. During both EEG recordings we presented “trained/matching” and “untrained/mismatching” pairings of multisensory stimuli. This distinction is only meaningful after the training of course, and comparison of data from the first versus the second EEG session provides a measure of the influence of learning of the pairs.

### Behavioral data analysis

For each individual training session we measured the percentage of correct responses. The learning curve was calculated on average data using an exponential nonlinear model to quantify the effect of learning over time, where time was defined by session number. The fit was performed using GraphPad Prism 8.0.1 with the following equation:

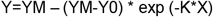

where Y0 is the intercept and YM is the maximum value (i.e. the plateau).

### Analysis of ERPs

EEG data were continuously recorded with a 128 channel Biosemi system and analyzed using EEGLAB 12_0_2_6b and ERPLAB 5.0.0.0 running under MATLAB 2014a (The Mathworks, Inc.). EEG recording was sampled at 512 Hz and a butterworth filter (0.1 Hz high-pass and 40 Hz low-pass) was applied to the continuous data. Bad channels were removed as were data containing large motion artifacts. Data were referenced to the average of the scalp channels.

The continuous EEG was segmented into epochs from 100 ms before to 500 ms after the onset of each stimulus. Baseline correction was performed by subtracting the mean value from the 100 ms before stimulus onset from the original signal. Large eye-movement artifacts (i.e. blinks/saccades) were identified and removed using an Independent Component Analysis (ICA). Bad channels were then interpolated and an artifact rejection with a voltage threshold of ±100 μV was applied. Individual event related potentials (ERPs) were calculated as the average of all the epochs for each experimental condition (unisensory visual, unisensory audio, audiovisual (and audiovisual match and mismatch for the post-training analysis). Target trials, which would include motor and other target related processes, were excluded from data analysis. ERPs were then averaged across participants into group averages for visualization and illustration purposes.

#### Testing the 3-stage model: The effect of learning on multisensory brain responses

To test the model’s predictions, and measure the effect of learning on multisensory brain responses, we compared the amplitude of the ERPs evoked by match/mismatch multisensory stimuli before and after training. For this assessment, we used separate repeated measure ANOVAs for each of the three temporal windows predicted by the model. For the first stage, we measured voltage between 40 to 90 ms post-stimulus-onset from six electrodes located over occipital scalp (Oz, O1, O2, Pz, P1 and P2). For the second stage, we expected multisensory object recognition to occur within sensory areas of the ventral pathway dedicated to object processing and recognition, like the LOC. For this reason we measured responses from three occipital and three frontal channels (the central Fz and the two lateral adjacent channels; channels were selected based on Molholm, Ritter, Javitt, & Foxe, 2004). The temporal window considered here is the one of the visual N1 (from 120 to 200 ms). For the third stage, we expected involvement of a widespread cortical network and we considered the same channels as in stage 2, but we analyzed data in a different temporal window (from 220 to 500 ms).

#### Exploratory Analyses: Hypothesis generation and a replication cohort

While initial analyses were strictly limited to the predictions of the 3-stage model, we also planned to explore the entire spatio-temporally rich data matrix, with the aim of discovering any potential missed effects due to the very limited/conservative a-priori analysis plan. In this second stage of analysis, if new/unpredicted effects were uncovered, we treated these outcomes as hypothesis generation tools, calculated the power needed to establish replicability of the effects, and conducted a direct replication study in a naïve cohort of participants. As will be discussed below, this resulted in the collection of data from an additional 10 participants and the following analyses were executed, including the full cohort (n = 30).

Object learning effects were assessed by comparing electrophysiological responses triggered by the presentation of multisensory matches and mismatches, before and after training. As mentioned before, we conducted a data-driven exploratory t-test analysis on the first 20 participants. Responses evoked from matches were subtracted from responses evoked by mismatches and then differences measured before training were compared with those measured after training, for all time points and all channels (see Supplemental Material for details about the analysis - https://figshare.com/articles/journal_contribution/SupplementalMaterial_NOPA_Foxe_doc/19172378). Effects were considered significant only for p-values <0.05 for at least 9 consecutive time points, i.e. 18 ms (Guthrie & Buchwald, 1991).

#### Quantifying multisensory integration: The general subadditive/superadditive effects of integration on brain responses

This last stage of analysis was performed on the full dataset of the 30 participants. General multisensory effects were measured as the difference between brain responses evoked by audiovisual stimuli and responses evoked by unisensory stimuli. To accomplish this we pooled data from the two audiovisual conditions (matches/mismatches) and subtracted the sum of the unisensory visual and audio responses (referred to as sum hereafter). We considered the following channels: Oz, O1, O2, Pz, P1,P2, Fz, F1 and F2. Temporal windows (80-100 ms; 130-150 ms; 250-500 ms) were selected based on previous literature and on the largest differences between the multisensory and the sum of the unisensory responses, within the expected time ranges.

## Results

Figure 2 shows the results of the behavioral training (for all the 30 participants). The exponential nonlinear model accurately captures the data (R =0.95) confirming that the number of correct responses increased exponentially with the training session. Indeed, as expected, all participants achieved nearly 100% correct responses by the completion of the training phase, and the plateau value (i.e. the critical point where the amount of correct responses can no longer increase) was 97.59. Notice that participants performed extremely well already in the second training session, and continued to improve up through the 5^th^ training session, then the performance stabilized. Based on this analysis, we are confident that all participants learned the associations between the new audiovisual objects, and we conjecture that subsequent training sessions served to more deeply encode these associations.

**Figure 2:**
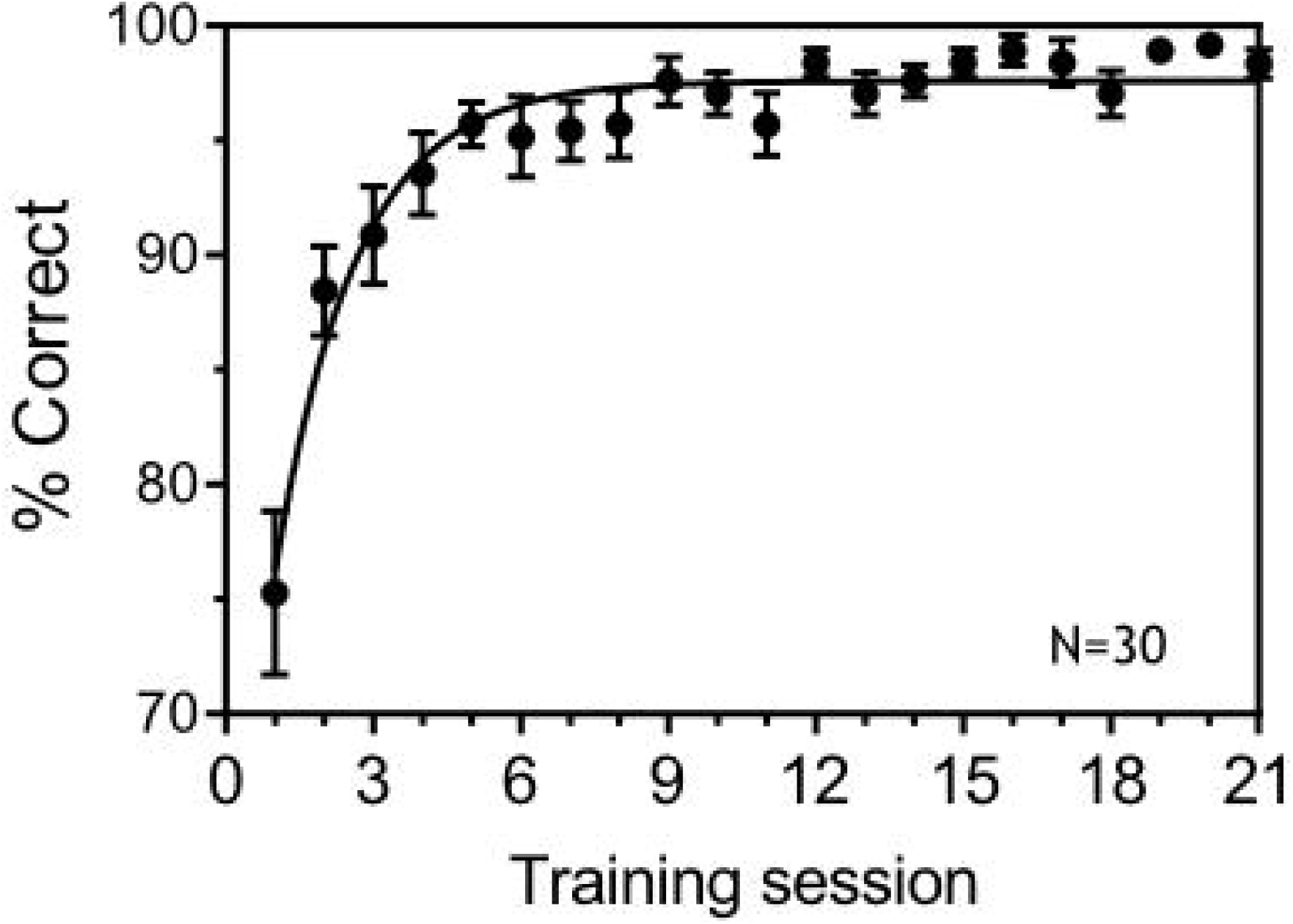
Behavioral performance during the training. The figure shows the percentage of correct responses as a function of the training session. Performance increases with time, and high accuracy is achieved by most of the participants already around the 6^th^ session (second day of training).

EEG data were first analyzed using a hypothesis-driven approach that considered the specific temporal windows predicted by the model. For the first stage of the model (40-90ms), we compared evoked potentials during matching and non-matching stimuli. As discussed in Methods, the analysis considered training (before/after), stimulus type (match/mismatch) and channels (Pz, P1, P2, Oz, O1, O2 – channel labels were renamed to match the 10-20 system that represents the official nomenclature) as factors. We found a significant effect of channel (F_(5, 95)_=22.31 P<0.0001 □^2^=0.54), stimulus type (F_(1, 19)_=12.19 P=0.002 □^2^=0.39) and a significant interaction between channel and stimulus type (F_(5, 95)_=9.19 P<0.0001 □^2^=0.32), but no effect of training (results from the analysis are reported in tables 1-3 of Supplemental Materials - https://figshare.com/articles/journal_contribution/SupplementalMaterial_NOPA_Foxe_doc/19172378). As planned, we repeated this same analysis for both the second and third temporal stages of the model. For the second stage, we considered the temporal window from 120 to 200 ms post-stimulus-onset, and the following channels: Fz, F1, F2, Oz, O1, O2. We found a significant effect of channel (F_(5, 95)_=6.39 P<0.0001 □^2^=0.25), stimulus type (F_(1, 19)_=15.10 P=0.001 □^2^=0.44), but again no effect of training and no interactions. For the third stage, we considered the temporal window from 220 to 500 ms post-stimulus-onset, and the same channels as stage two. Here, we only found a significant effect of channel (F_(5, 95)_=110.40 P<0.0001 □^2^=0.85). This first analysis did not reveal any learning effect induced by the training, within these specific temporal windows predicted by the model, and also suggests differences in the electric potentials evoked by matching and non-matching multisensory stimuli, even before the training, when participants had not yet been informed of any basis upon which to discriminate between the two categories (i.e. the match mismatch distinction only becomes germane when the participants begin the training sessions, and during the initial EEH session, they are entirely naïve to nay potential future relationships between the audio-visual elements). Results also suggest differences in the potentials recorded across different channels.

In the second exploratory analysis stage, we embraced a data-driven approach to better identify the most influential temporal windows where learning effects occurred. In the first 20 participants, significant effects were found in the temporal windows between 100 and 150 and between 250 and 300 ms poststimulus onset. For the first temporal window (100-150 ms), learning effects were located on a central cluster centered on the Cz channel (clusters are reported in Supplemental Figure S5 of Supplemental Material - https://figshare.com/articles/journal_contribution/SupplementalMaterial_NOPA_Foxe_doc/19172378). The multiple t-test analysis was followed up with two repeated measure ANOVAs, with temporal windows and channels of analysis defined by the above data-driven analysis. The analysis of the 100 to 150 ms window comprised the factors: channels (10 channels defining the cluster), training (before/after) and stimulus type (match/mismatch) and revealed a significant effect of channel (F_(5,145)_=8.32; p<0.0001; □^2^=0.22), training (F_(1,29)_=12.70; p=0.001; □^2^=0.30), and stimulus type (F_(1,29)_=27.30; p<0.0001; □^2^=0.48), and significant interactions between channel and training (F_(5,145)_=22.07; p<0.0001; □^2^=0.43), channel and stimulus type (F_(5,145)_=33.52; p<0.0001; □^2^=0.53), and channel, training and stimulus type (F_(5,145)_=48.56; p<0.0001; □^2^=0.62). For the second temporal window (250-300 ms), learning effects were focused over centro-parietal scalp (Pz). We considered as factors: channels (the 6 central channels defining the cluster), training (before/after) and stimulus type (match/mismatch) and found a significant effect of channel (F_(9,171)_=10.16; p<0.0001; □^2^=0.34), and significant interactions between channel and training (F_(9,171)_=57.86; p<0.0001; □^2^=0.75), channel and stimulus type (F_(9,171)_=10.76; p<0.0001; □^2^=0.36) and channel, training and stimulus type (F_(9,171)_=30.87; p<0.0001; □^2^=0.62). We defined these clusters as Regions of Interest (ROIs) and then collected data from 10 more participants (sample size was determined on the basis of a Power analysis grounded on our previous analysis, see Supplemental Material - https://figshare.com/articles/journal_contribution/SupplementalMaterial_NOPA_Foxe_doc/19172378) and measured learning effects in our ROIs to determine if these “discovered” effects were replicable. When replicability was established (see Supplemental Material - https://figshare.com/articles/journal_contribution/SupplementalMaterial_NOPA_Foxe_doc/19172378), we pooled data across all 30 participants for further analyses.

EEG data related to early multisensory effects (80-100ms post stimulus presentation; n=30) are displayed in figure 3. The left panel shows EEG results before training, whereas the right panel of the figure illustrates post-training data. Activity from three occipital channels (left, right and center) and from the three parietal channels (left, right and center) are shown and the temporal window is highlighted with a yellow shade in all the graphs. In each case data from both the multisensory condition (pink line) and for the sum of the two unisensory conditions (auditory+visual; blue line) are plotted. Multisensory effects are evident as early as 80 ms post-stimulus-onset and, in line with the model, such early effects are visible even before training. To test for significant differences across conditions, we ran a repeated measures ANOVA with factors: channels (A5, A19, A32, A15, A23, A28 – channels label correspond to the 128channels Biosemi system where A23=Oz and A19=Pz), training (before/after) stimulus type (multisensory/unisensory sum). We found a significant effect of channel (F_(5, 145)_=9.44 P<0.0001 □^2^=0.25), stimulus type (F_(1, 29)_=35.45 P<0.0001 □^2^=0.55), channel and training (F_(5,145)_=18.42 P<0.0001 □^2^=0.38) channel and stimulus (F_(5, 145)_=39.42 P<0.0001 □^2^=0.57) and channel, training and stimulus (F_(5, 145)_=43.87 P<0.0001 □^2^=0.60). The posterior distribution of this early multisensory effect is highlighted in topographic maps for pre and post training sessions.

**Figure 3:**
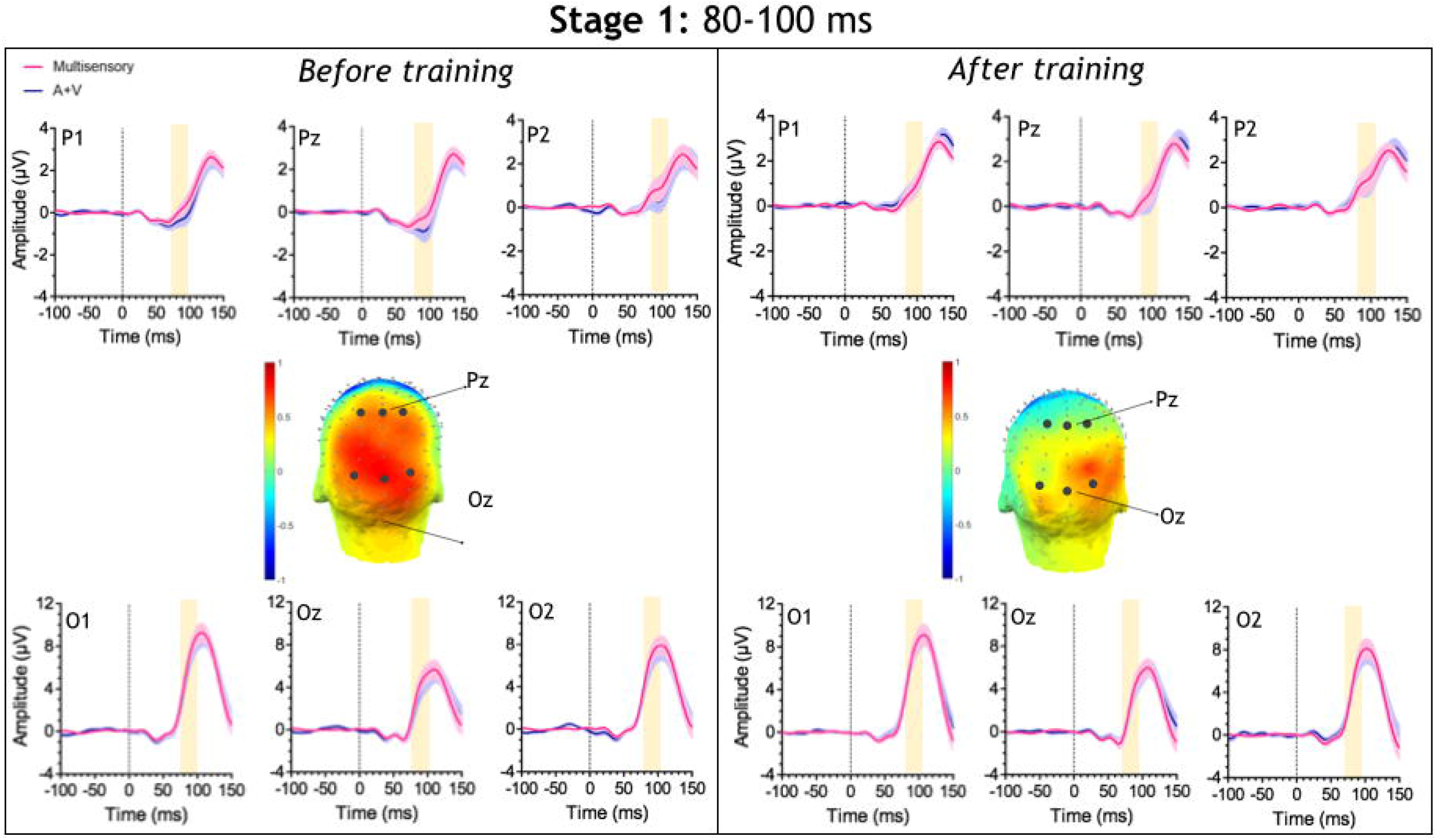
Data for the first stage of the model (80-100 ms post-stimulus onset) before (left panel) and after (right panel) the training. The two scalp maps show the mathematical difference between the multisensory condition and the sum of the two uinisensory conditions. The upper graphs show the voltage for three parietal electrodes (note that channel A19 in the 128-channel biosemi system corresponds to the Pz channel of the 10-20 system), and the lower graphs show the voltage for 3 occipital channels (channel A23 = Oz). The pink line represents the multisensory condition and the purple line the unisensory sum condition (A+V). The yellow shade simply highlights the temporal window that we considered for the analysis.

Figure 4 shows the multisensory and sum ERPs over occipital and frontal scalp for pre- and posttraining phases (left and right sides of the figure, respectively). Multisensory response is larger (i.e. more negative in occipital channels and more positive in frontal channels) than the sum response in the 130 to 150 ms timeframe, corresponding to the second stage of the model. Topographical maps of the multisensory minus sum response illustrates the posterior and frontal topography of this difference, and that it becomes more pronounced following training. In this second temporal window (highlighted by a green shaded bar in all the graphs), we found a significant effect of channel (F_(5, 145)_=4.33 P=0.001 □^2^=0.13), training (F_(1, 29)_=4.62 P=0.04 □^2^=0.13), stimulus type (F_(1, 29)_=4.17 P=0.05 □^2^=0.12), channel and training (F_(5,145)_=3.85 P=0.003 □^2^=0.12) channel and stimulus (F_(5, 145)_=39.42 P=<0.0001 □^2^=0.57) and training and stimulus (F_(1,29)_=14.01 P=0.001 □^2^=0.32).

**Figure 4:**
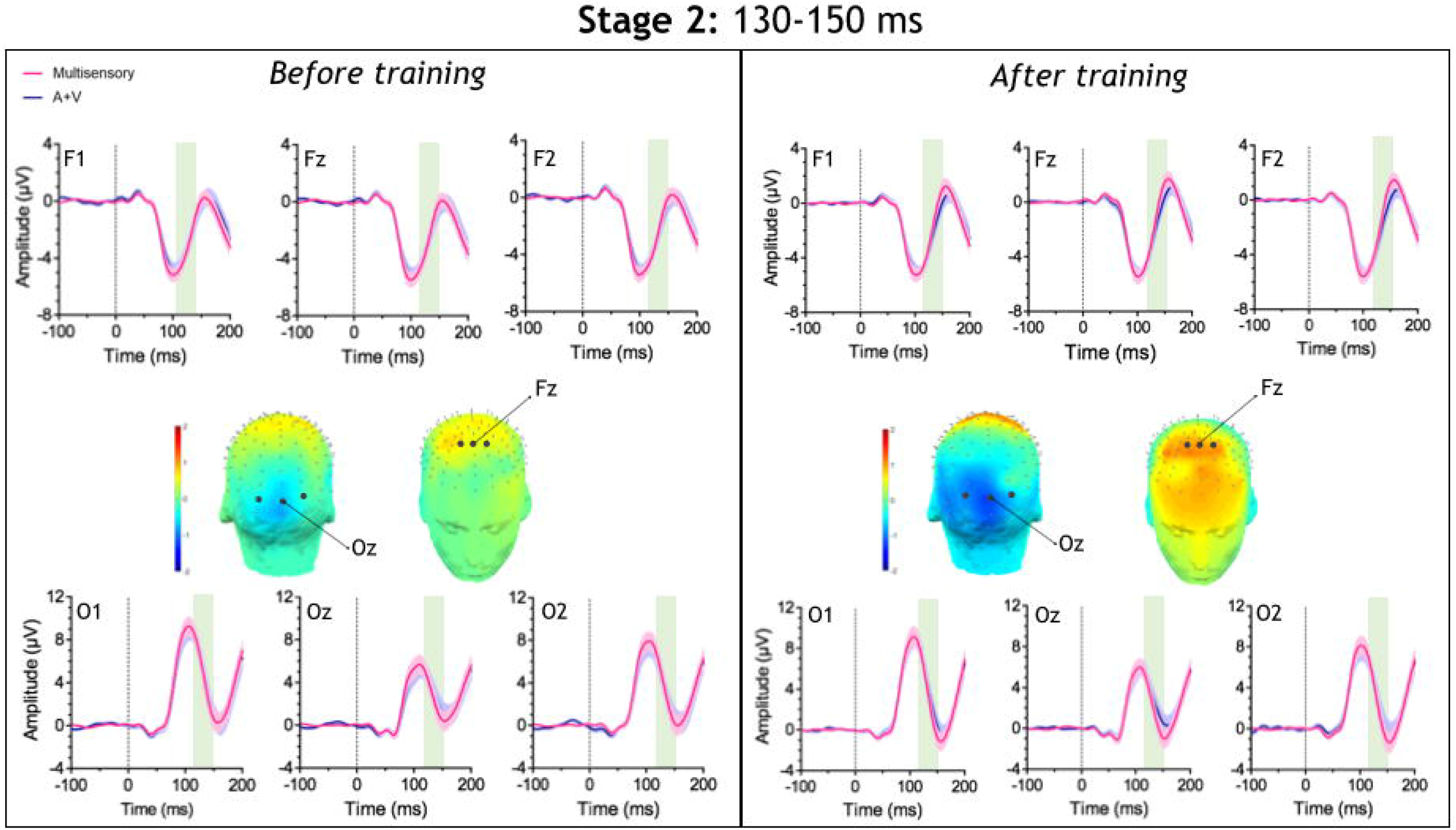
Data for the second stage of the model (130-150 ms post-stimulus onset) before (left panel) and after (right panel) the training. The two scalp maps in the upper boxes show the difference between the multisensory and the unisensory sum condition and the graphs contained in the same boxes show the voltage for 3 occipital channels (lower panels, channel A23 = Oz) and for 3 frontal channels (higher panels, channel C21 = Fz). The pink line represents the multisensory condition and the purple line the unisensory sum condition (A+V). The green shade simply highlights the temporal window that we considered for the analysis.

We further analyzed the effects of the behavioral training by comparing electrophysiological responses evoked by match versus mismatch audiovisual (AV) pairs, and results are illustrated in figure 5. As previously explained in the EEG analysis section, here we considered a cluster of central channels and a temporal window between 100 and 150 ms post-stimulus onset, which is close to both the auditory N1 (~100 ms) and the visual N1 (~150 ms) latency and to the time frame that we were considering for stage 2 of the model. We ran a repeated measure ANOVA with factors: channel (the 14 channel composing the cluster), training (before/after) and stimulus type (match/mismatch) and found significant effects of channel (F_(13, 377)_=39.59 P<0.0001 □^2^=0.57), training (F_(1, 29)_=16 P<0.0001 □^2^=0.36), and stimulus type (F_(1, 29)_=23.36 P<0.0001 □^2^=0.44), and significant interactions between channel and training (F_(13,377)_=35.83 P<0.0001 □^2^=0.55), channel and stimulus type (F_(13,377)_=56.82 P<0.0001 □^2^=0.66) and channel, training and stimulus type (F_(13,377)_=24 P<0.0001 □^2^=0.46). Figure 5 shows topographical maps of the difference in the electrical activity evoked by match and mismatch stimuli (match-mismatch) before and after training. The graphs display the average voltage recorded in the channels that compose the cluster. These data confirm that learning affects the magnitude of electrical responses triggered by multisensory object pairings.

**Figure 5:**
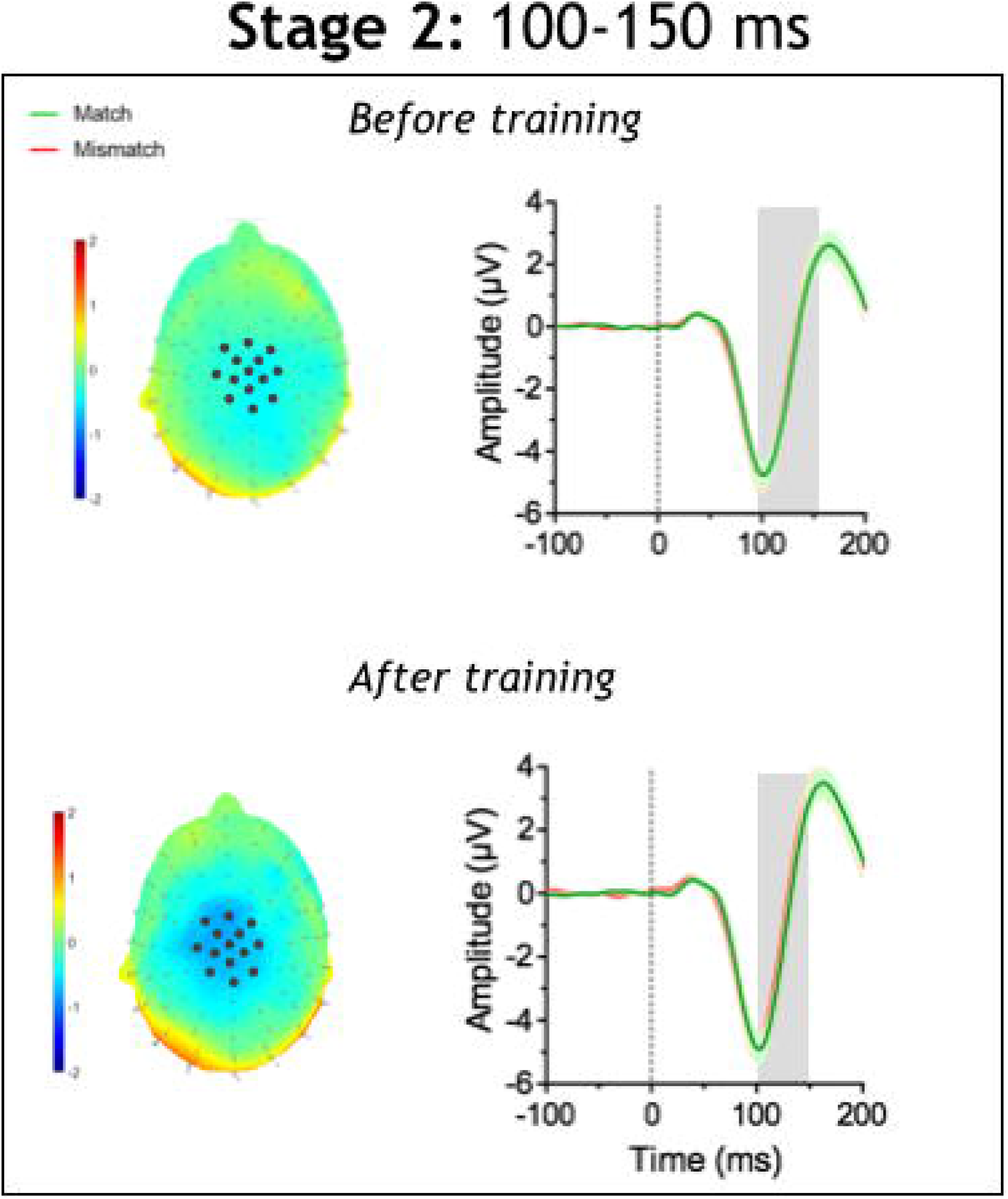
The scalp maps show the differences between the match and the mismatch condition. Similarly, the report the average voltage, calculated over all the channels that constitute the ROI, to match (green line) and mismatch (red line) stimuli. Here we found significant differences in the temporal window between 100 to 150 ms (the temporal window is highlighted by the grey shade).

The results for the third time-window, corresponding to the third stage of the model, are presented in figure 6. The topographic maps suggest that late multisensory interactions take place in a widespread network of brain areas. In this temporal frame, electrophysiological responses by multisensory stimuli are more negative than those evoked by the sum of unisensory responses recorded in the occipital channels, and more positive in the frontal channels. The difference between the two conditions is highlighted by the scalp maps and the waveforms showing activity recorded in the multisensory and the summed unisensory conditions for the 3 occipital (bottom graphs) and 3 frontal (upper graphs) channels. In this last temporal window (highlighted with a purple shade in all the graphs), we found a significant effect of channel (F_(5,145)_=94.73 P<0.0001 □^2^=0.76), of stimulus type (F_(1, 29)_=136.71 P<0.0001 □^2^=0.82), and a significant interaction between channel and training (F_(5,145)_=108.46 P<0.0001 □^2^=0.78), channel and stimulus type (F_(5,145)_=92.11 P<0.0001 □^2^=0.76), training and stimulus type (F_(1,29)_=47.66 P<0.0001 □^2^=0.62) and between channel, training and stimulus type (F_(5,145)_=101.69 P<0.0001 □^2^=0.77).

**Figure 6:**
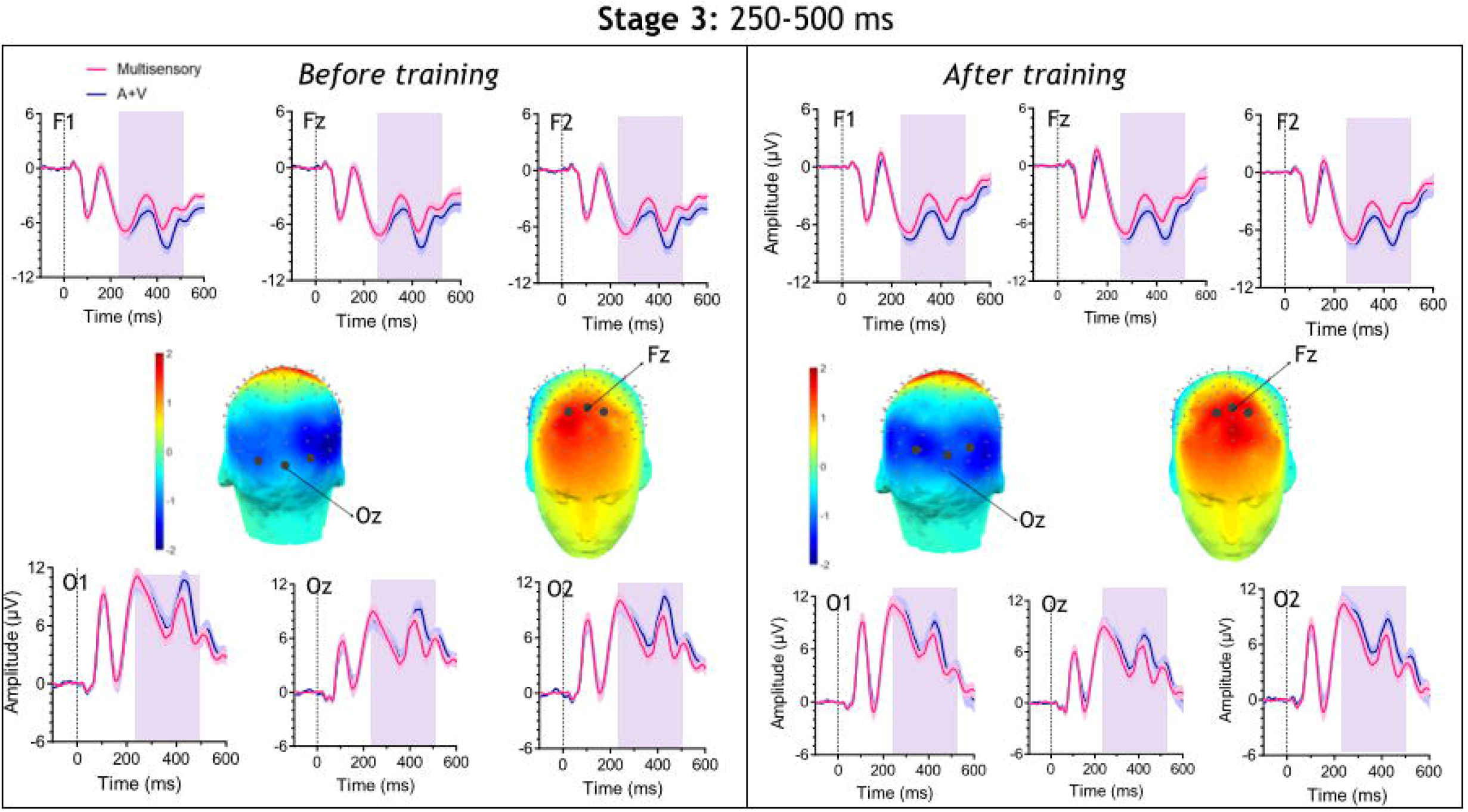
Data for the third stage of the model (250-500 ms post-stimulus onset) before (left panel) and after (right panel) the training. The two scalp maps show the difference between the multisensory condition and the unisensory sum condition. The upper graphs show the voltage for three parietal electrodes (channel A19 = Pz), and the lower graphs show the voltage for 3 occipital channels (channel A23 = Oz). The pink line represents the multisensory condition and the purple line the unisensory sum condition (A+V). The purple shade highlights the temporal window that we considered for the analysis.

Also for the third stage, we assessed learning effects induced by the behavioral training. Specifically, we compared electrical responses evoked by multisensory stimuli that were matches to those evoked by multisensory stimuli that were mismatches, before and after the training in the temporal window between 250 and 300 ms post-stimulus onset. For this analysis we consider a second cluster of channels that was located on the parietal region of the scalp (see EEG analysis and Supplemental Material for details - https://figshare.com/articles/journal_contribution/SupplementalMaterial_NOPA_Foxe_doc/19172378). Figure 7 shows difference in the electrical activity evoked by match and mismatch stimuli (match-mismatch) before and after the training. The left panel of the figure displays topographical maps of the average difference in the electrical activity recorded in the temporal window of interest, while the graphs display the time course of the average voltage recorded in the channels that compose our second ROI. Here, the repeated measure ANOVA revealed significant effects of channel (F_(9,261)_=11.17 P<0.0001 □^2^=0.27), of stimulus type (F_(1, 29)_=5.43 P=0.03 □^2^=0.15), and a significant interaction between channel and training (F_(9,261)_=50.68 P<0.0001 □^2^=0.63), channel and stimulus type (F_(9,261)_=14.34 P<0.0001 □^2^=0.33), and between channel, training and stimulus type (F_(9,261)_=38.63 P<0.0001 □^2^=0.57).

**Figure 7:**
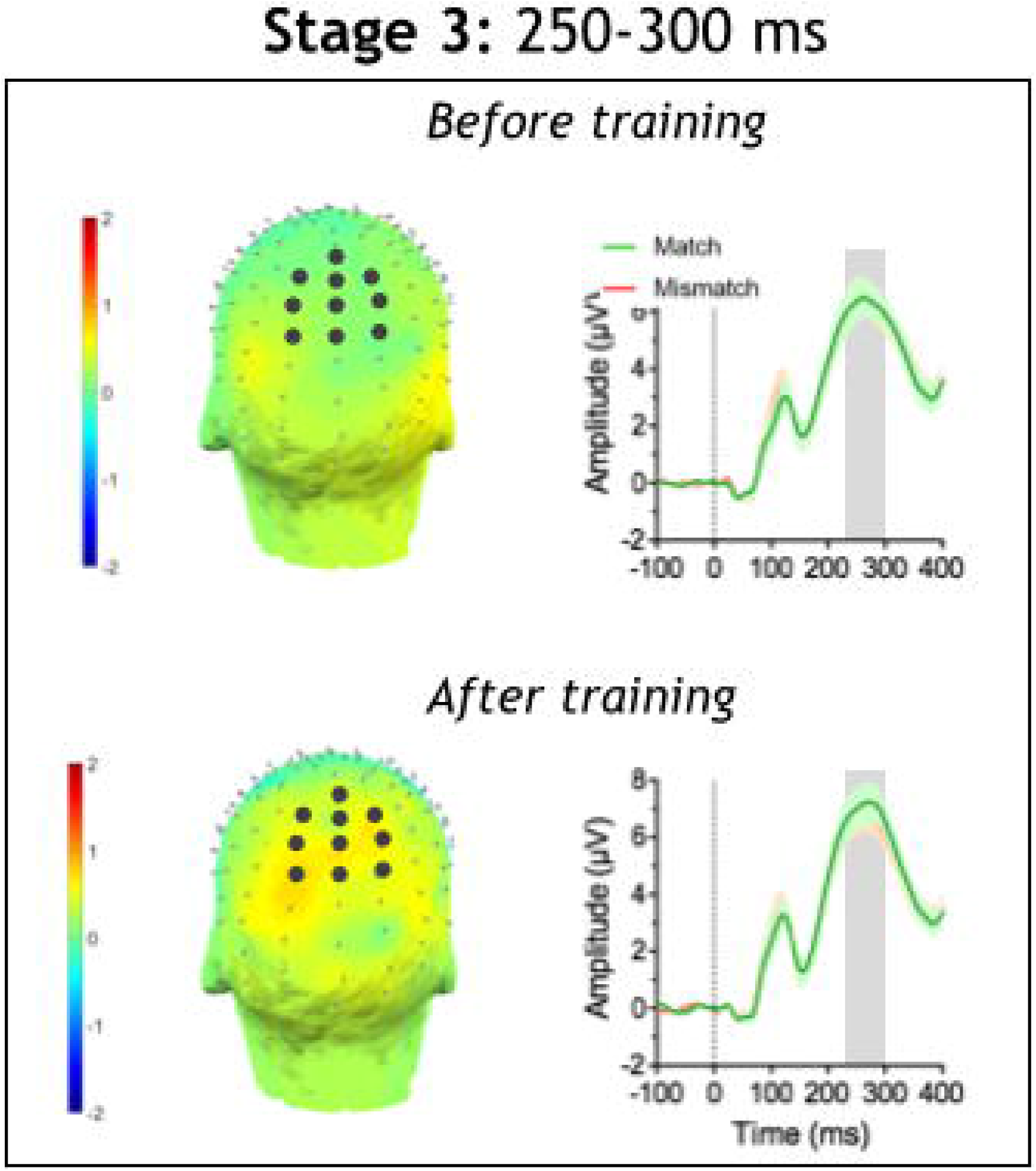
Differences between responses evoked by match and mismatch condition stimuli. The graphs report the average voltage, calculated over all the channels that constitute the ROI, to match (green line) and mismatch (red line) stimuli. Here we found significant differences in the temporal window between 100 to 150 ms (the temporal window is highlighted by the grey shade).

## Discussion

How do multisensory representations of objects change over time as multisensory objects become part of our lexicon? Here we had participants learn a set of novel audio-visual objects, to examine how their neural representations would change as they went from being randomly presented co-occurring stimuli to systematically paired recognizable multisensory objects. We formulated a rudimentary three stage model, based on observations from the literature, from which to generate predictions about the spatio-temporal dynamics of multisensory object representation and how this would change as a function of training.

Many studies have shown that auditory and visual inputs are integrated early in the temporal (<100 ms) and anatomical (primary or secondary sensory cortices) cortical processing hierarchy (Cappe, Thut, Romei, & Murray, 2010; John J. Foxe et al., 2000; Giard & Peronnet, 1999; Molholm et al., 2002; Murray et al., 2005; Raij et al., 2010; Shams, Iwaki, Chawla, & Bhattacharya, 2005; Mark T. Wallace, Meredith, & Stein, 1998). We assumed that these early interactions would be present for all our stimuli, and that, being so early, they would not modulate due to training with specific audio-visual pairs. In agreement with these predictions, our results showed that brain electrical responses evoked by novel multisensory objects were different than the sum of the unisensory responses at early latencies (80-100ms), and that these multisensory effects were consistent with generators in sensory cortices. What is more, while there was an effect of training such that the MSI effect decreased after training, this did not interact with audiovisual congruency. In other words, after training, the early MSI effect did not differ for learned versus new multisensory pairings. Because learning was not a determining factor at this first stage of multisensory processing in our data, we suggest that this early integration serves as a simple detection system (Schroeder & Foxe, 2002) to establish multisensory binding in the next stages of processing. These early multisensory effects might be crucial for integration at later stages of processing, specifically in determining whether multiple sensory signals belong to the same source or need to be treated as separate objects. Future studies should validate this hypothesis.

The generalized effect of exposure to the stimuli during training, an effect that we did not anticipate, showed reduced multisensory responses that were more lateralized to the right hemisphere (see figure 3, right panel). One possible explanation is that novel stimuli trigger attention-related activation (Escera, Alho, Winkler, & Näätänen, 1998), resulting in larger and more widespread brain activity. Another possibility is that repetition effects (Kalanit Grill-Spector, Henson, & Martin, 2006) are at play, leading to a diminution of the sensory responses, which is in turn compounded in the multisensory responses.

We predicted a second stage of multisensory object processing that would modulate as a function of learning of the audiovisual pairs, in the 100 to 200 ms timeframe and over lateral occipital regions. The visual N1 is an electrophysiological response that peaks at about 170ms and has been closely associated with object processing (Bar, 2003; Clark, Fan, & Hillyard, 1994; Di Russo, Martínez, Sereno, Pitzalis, & Hillyard, 2002; Molholm et al., 2004). Generators of the visual N1 have been typically identified within perceptual association regions of cortex, like the LOC, which is known to be involved in the representation of congruent sounds and images (A. Amedi et al., 2005; Besle, Fort, Delpuech, & Giard, 2004; Murray et al., 2004). Prior brain imaging work also supports the idea that LOC processes multisensory object representations, not only for audiovisual stimuli, but also across the visual and haptic modalities (Amir Amedi, Malach, Hendler, Peled, & Zohary, 2001; Lucan et al., 2010). Specifically, LOC appears to represent more abstract “amodal” stimulus properties, like object category, that are accessible through multiple sensory modalities (A. Amedi et al., 2005; Erdogan et al., 2016; K Grill-Spector & Malach, 2001; Kourtzi, Erb, Grodd, & Bülthoff, 2003; Lacey, Tal, Amedi, & Sathian, 2009; Stilla & Sathian, 2008). Consistent with our predictions, we found differences between multisensory responses (i.e. match + mismatch stimuli) and the summation of the auditory and the visual responses. These multisensory effects were focused over occipital and lateral occipital (left side) scalp areas as well as over more frontal scalp regions (see figure 4). Critically, we also found post-training changes in the electrophysiological signal evoked by the stimuli that were learned during training (i.e. matches) in this same timeframe, although with a topography over centro-parietal scalp. Even though highly speculative at this point, this could reflect generators in the temporal pole, which is a convergence region associated with multimodal conceptual representation (Olson, Plotzker, & Ezzyat, 2007; Skipper, Ross, & Olson, 2011). This second stage of processing, in which visual object recognition areas were engaged by multisensory object learning, is in line with the notion that higher-order regions of the cortex are characterized by their functionality rather than by the sensory information that they process (J. John Foxe & Molholm, 2009; Lucan et al., 2010; Pascual-Leone & Hamilton, 2001).

It is important to consider that once multisensory objects are established through learning, presentation of just one of the sensory elements is likely to lead to activation of the entire multisensory network (Lacey, Flueckiger, Stilla, Lava, & Sathian, 2010; Lucan et al., 2010; Von Kriegstein & Giraud, 2006; Von Kriegstein, Kleinschmidt, Sterzer, & Giraud, 2005). As such, traditional tests of MSI comparing the summed unisensory responses to the multisensory response might underestimate the strength of multisensory effects. Moreover, sensory stimuli that are semantically related when presented singularly can induce a top-down spread of attention, activating a preexisting cortical representation of the well-known multisensory object (Fiebelkorn et al., 2010). Indeed, for this particular stage, we found a general effect of training, suggesting that even unisensory responses might have been modulated by the learning process.

Finally, we predicted that a third stage of multisensory object processing, in a temporal window typically associated with conceptual or cognitive-level processing (250 to 500ms), would be engaged by the presentation of novel but not well-learned multisensory objects. That is, we predicted that before training, when the auditory and the visual information have no known relationship to each other, they will be further analyzed for potential binding. However, after multisensory associations are learned, stage-3 processing should become unnecessary and subsequently diminish as stage-2 processing progresses. Instead what we found is that in this later timeframe, multisensory effects increased after training, but that this did not differ for learned versus novel pairings.

On the other hand, in a post hoc analysis we identified multisensory responses related to object recognition within a temporal window (250-300 ms) included in our rudimentary stage 3. Here the response was bigger for learned multisensory pairs, and focused over parietal regions. In this temporal window, a modulation of multisensory responses induced by object recognition (trained versus untrained pairs) might interact with memory processes. Further research can thus shed light on the dynamics and function of this late stage of multisensory processing.

Tracking down the neurophysiological markers of multisensory learning was the primary goal of this research. Specifically we aimed to identify and distinguish multisensory responses that occur: when the brain is initially exposed to novel multisensory stimuli; when the unisensory components of these stimuli start to be associated as a result of repeated exposure; when multisensory representations are consolidated and the brain can identify them as familiar objects. Surprisingly and contrary to our expectations, multisensory effects related to learning were modest in size and distributed over large regions of the scalp, leading to the general conclusion that there is no neurophysiological marker that can be clearly associated to multisensory object recognition. Of course we may speculate that our paradigm has several limitations as the behavioral training wasn’t particularly challenging and learning was rather rapid. In real life, we learn to identify new multisensory objects after numerous exposures. New memories (such as new multisensory representations) might consolidate slowly over time and learning mechanisms could differ as a function of exposure. Moreover, because age is a crucial aspect of experience-dependent cortical plasticity (Knudsen, 2004) with young brains being more responsive to experience than older brains, deeper sensory learning in adults may require a more extended period of training. As a follow up study, it would be interesting to train participants with a more effective paradigm that involves the exposure to novel multisensory objects over a larger temporal interval. Another interesting explanation is that adults may assimilate new sensory experiences into pre-existing consolidated knowledge and that the encoding, consolidation, and retrieval of new multisensory associations may therefore be influenced by long-term stored representations of the external world. This hypothesis could be possibly verified with a follow-up study in a younger population of children.

Research on multisensory integration in humans has radically increased in recent years, yet many fundamental questions remain to be explored. Understanding the “normal” mechanisms of multisensory integration is of key clinical relevance to assess and identify several neurological disorders that have been associated with deficits in MSI, like autism (Brandwein et al., 2013), schizophrenia (Ross et al., 2007), dyslexia (Hahn, Foxe, & Molholm, 2014), and many others. In this study, we characterized the temporal dynamics of audiovisual object processing and were able to identify multiple cortical stages at which multisensory interactions occur.

## Funding

Initial pilot funding for this work was provided by a grant from the Schmitt Program in Integrative Neuroscience (SPIN), through the University of Rochester’s Del Monte Institute for Neuroscience. Recruitment and phenotyping at The Cognitive Neurophysiology Laboratory is conducted in collaboration with cores of the University of Rochester Intellectual and Developmental Disabilities Research Center (UR-IDDRC), which is funded by a center grant from the Eunice Kennedy Shriver National Institute of Child Health and Human Development (NICHD P50 HD103536 – to JJF).

## Notes

https://figshare.com/articles/journal_contribution/SupplementalMaterial_NOPA_Foxe_doc/19172378

### Competing Interest Statement

The authors have declared no competing interest.

### Summary of Updates

To fix broken links to supplemental materials

