## Supplemental Materials for "Learning a new class of multisensory associations: High-density electrophysiological mapping of the temporal course of audio-visual object processing" (from another version of this preprint: version 1, 2021-11-15)

### **Methods and procedures**

Visual and auditory stimuli adopted in the current study are reported below in Supplemental Figure S1 and S2. During the EEG recording, participants were fixating at a yellow cross that was presented at the center of the screen, while stimuli were delivered with an Inter-Stimulus-Interval ranging between 800 to 1500 ms. Stimuli could have been unisensory auditory/visual, multisensory match/mismatch, or target trials. Visual stimuli were displayed centrally on a grey background. The duration of all the stimuli was 300 ms. Multisensory stimuli consisted of the simultaneous presentation of a visual and an auditory stimulus. Target trials were similar to multisensory trials, but the auditory or the visual stimulus was replaced by auditory (6 trials) or visual (6 trials) noise. For the behavioral task, participants were asked to respond to target trials with a button press using their right index finger.

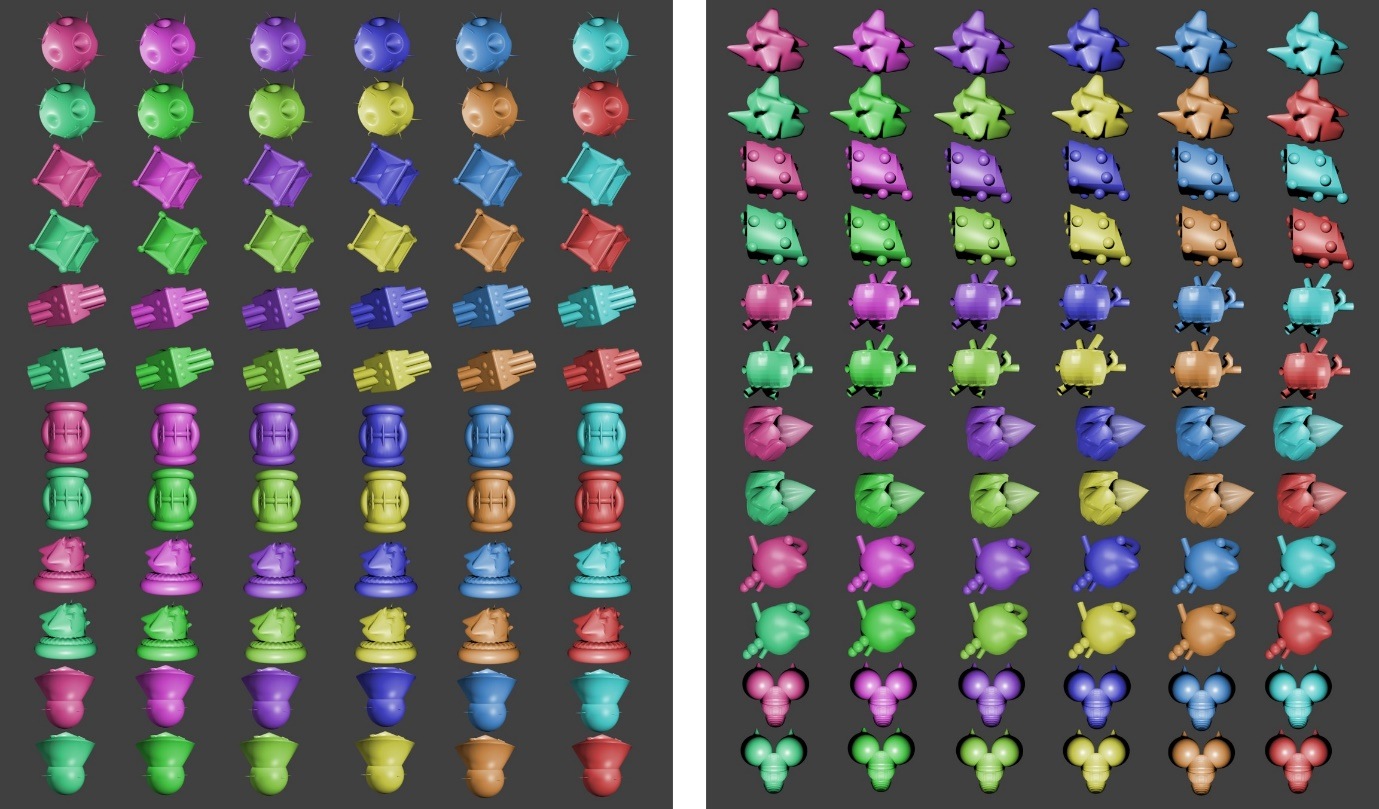

Supplemental Figure S1: List of all the visual stimuli adopted during the experiment (12 shapes combined with 12 colors)

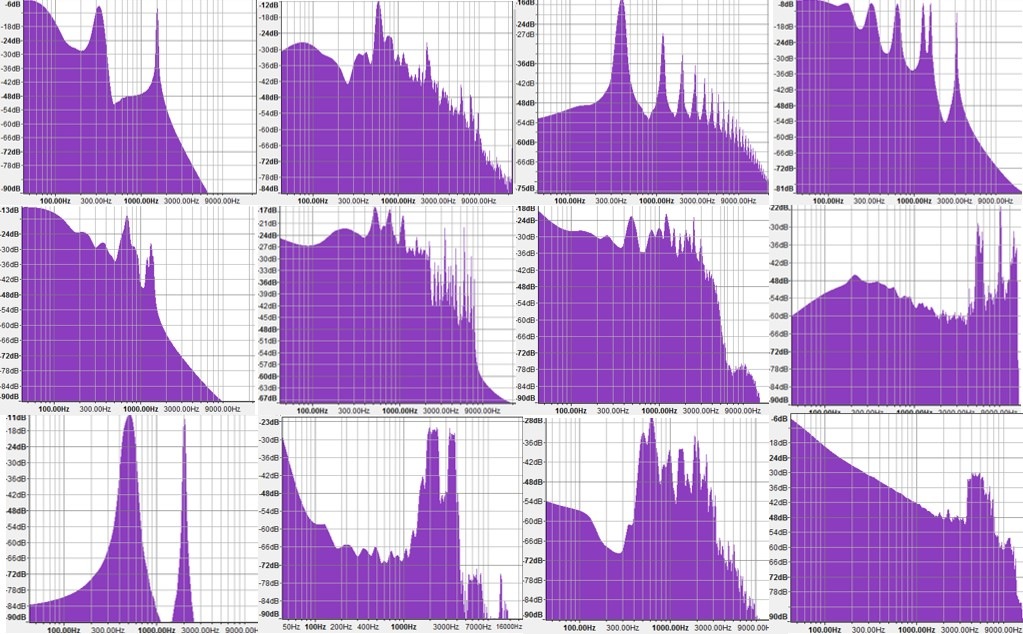

Supplemental Figure S2: Spectrograms of the 12 sounds created for the experiment

The training was performed through a touch screen tablet. For the exploration, the app displayed a grid of 12 blue circles, one for each multisensory object (see figure S4). After the selection of one of the circles, the grid disappeared and one of the multisensory stimuli was displayed for 300 ms. The selection of the same circle triggered the presentation of the same multisensory stimulus, but the color of the visual shape was always randomized. The spatial organization of the multisensory objects within the grid varied across sessions, but the multisensory pairings remained the same. For each session, participants were allowed to 36 exploration of the multisensory objects (on average 3 views per object).

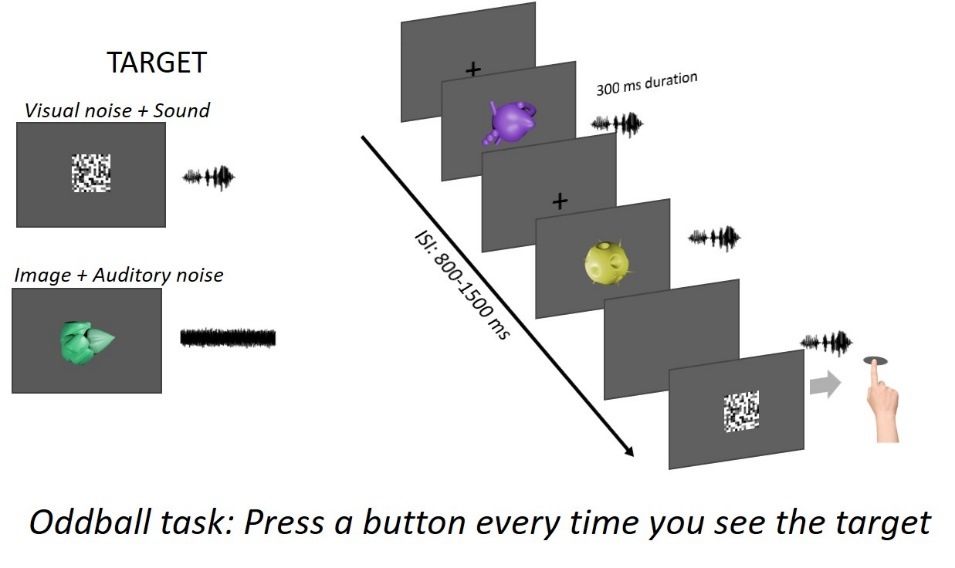

Supplemental Figure S3: Stimulus presentation procedure during the EEG session.

The test was a 2AFC task involving the presentation of 2 stimuli: a match, i.e. one of the multisensory object learnt during the exploration phase, and a mismatch, i.e. the wrong association between a visual shape and a sound (see figure S4). Participants were allowed to explore the stimuli four times (on average 2 views per stimulus) and were then required to identify the match. At the end of each training session, the software provided a feedback conveying information about the number of correct responses.

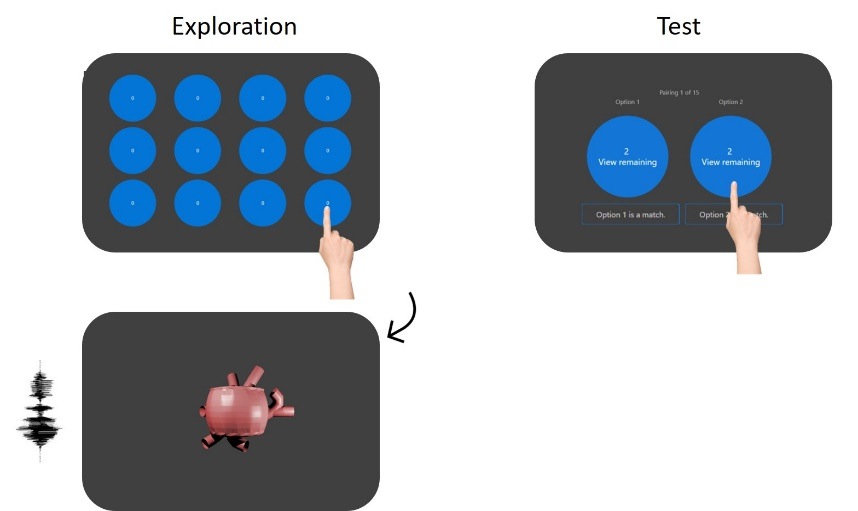

Supplemental Figure S4: Training procedure during exploration and test.

### **Testing the model:**

Table 1-3 show the results of the hypothesis driven analysis to test the model. Three repeated measure ANOVAs were performed for each of the three temporal windows predicted by the model. Factors were: channel (Oz, O1, O2, Pz, P1 and P2), stimulus type (match/mismatch) and training (before /after).

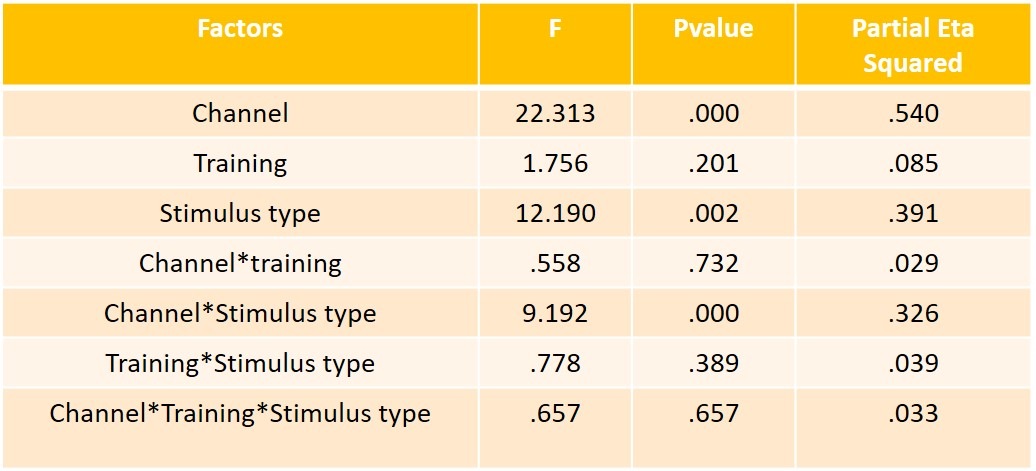

Table 1: Stage 1, matches Vs. mismatches. Comparison between the mean amplitude in the temporal window predicted by the model (40-90 ms). Analysis performed on the first 20 participants.

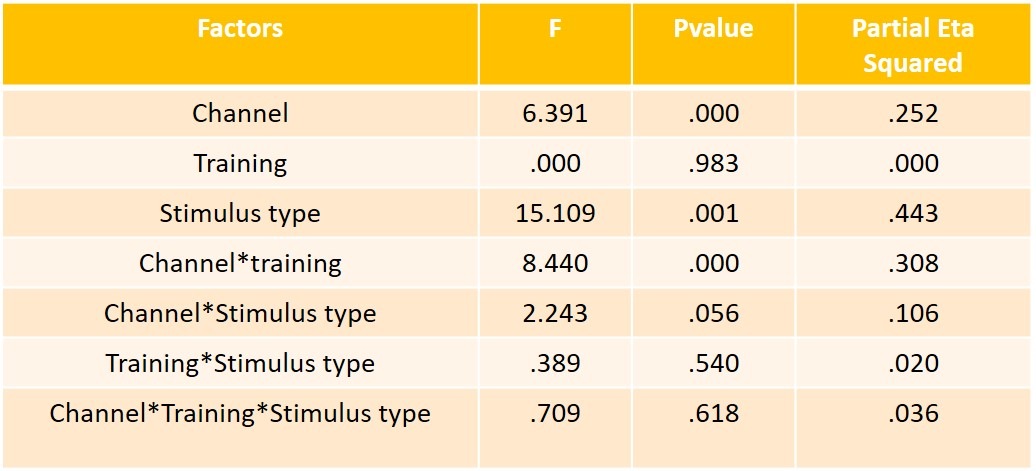

Table 2: Stage 2, matches Vs. mismatches. Comparison between the mean amplitude in the temporal window predicted by the model (120-200 ms). Analysis performed on the first 20 participants.

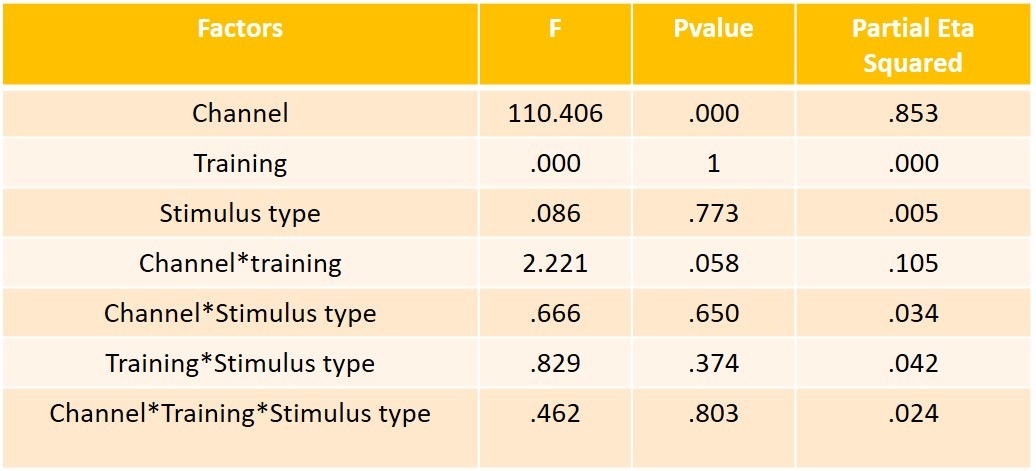

Table 3: Stage 3, matches Vs. mismatches. Comparison between the mean amplitude in the temporal window predicted by the model (220-500 ms). Analysis performed on the first 20 participants.

### **Exploratory analyses**

We conducted a data-driven exploratory t-test analysis on the first 20 participants to assess the effect of learning on multisensory responses. Learning effects were measured by subtracting the responses evoked by matches from the responses evoked by mismatches and then by comparing those responses before and after the training, for all time points and all channels. Only for visualization purpose of the statistical results (see figure S6), we grouped channels based on spatial proximity, scalp location and hemispheric differences in sensory processing (figure S5). We selected 3 frontal clusters (left, right and central – highlighted in the figures by the red lines), 3 central clusters (left, right and central – highlighted in the figures by the blue lines), 3 parietal clusters (left, right and central – highlighted in the figures by the black lines), 3 occipital clusters (left, right and central – highlighted in the figures by the green lines) and 2 temporal clusters (left and right – highlighted in the figures by the yellow lines).

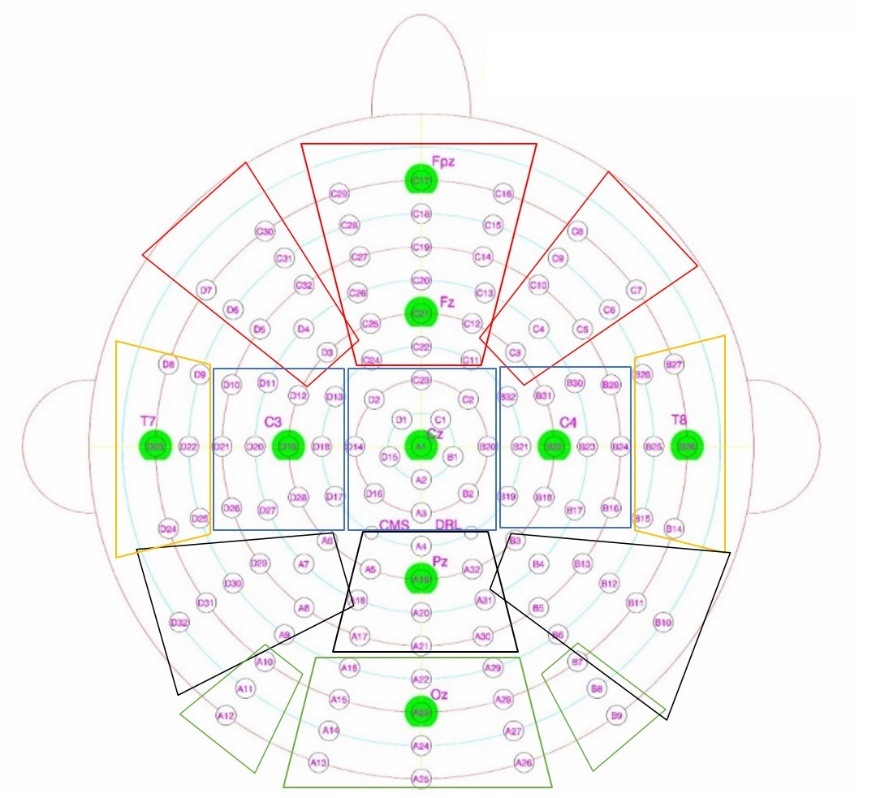

Supplemental Figure S5: Channel organization into multiple clusters.

For every channel within each cluster we subtracted the signal evoked by mismatch stimuli from that one evoked by match stimuli and compared these differences before and after training, for each time points. In figure S6 we reported t-values only for those comparisons that were significant (i.e. p<0.05) for at least 9 consecutive time points (i.e. 18 ms). Each line represents a channel and channels are grouped into clusters (see figure S5). We identified two significant modulations induced by learning: the first one appeared in a temporal window between 100 to 150 ms post-stimulus onset and was localized in a central cluster; and the second one localized in a central parietal cluster from 250 to 300 ms post-stimulus onset.

We conducted two repeated measure ANOVAs, one for each temporal window, to determine the significant of these learning effects. This analysis was first performed on 20 participants. For the 100-150 ms window we considered the following factors: channels (10 channels defining the cluster), training (before/after) and stimulus type (match/mismatch). Similarly, for the second temporal window we considered the following factors: channels (6 channels defining the cluster), training (before/after) and stimulus type (match/mismatch). Results are reported below in tables 4.

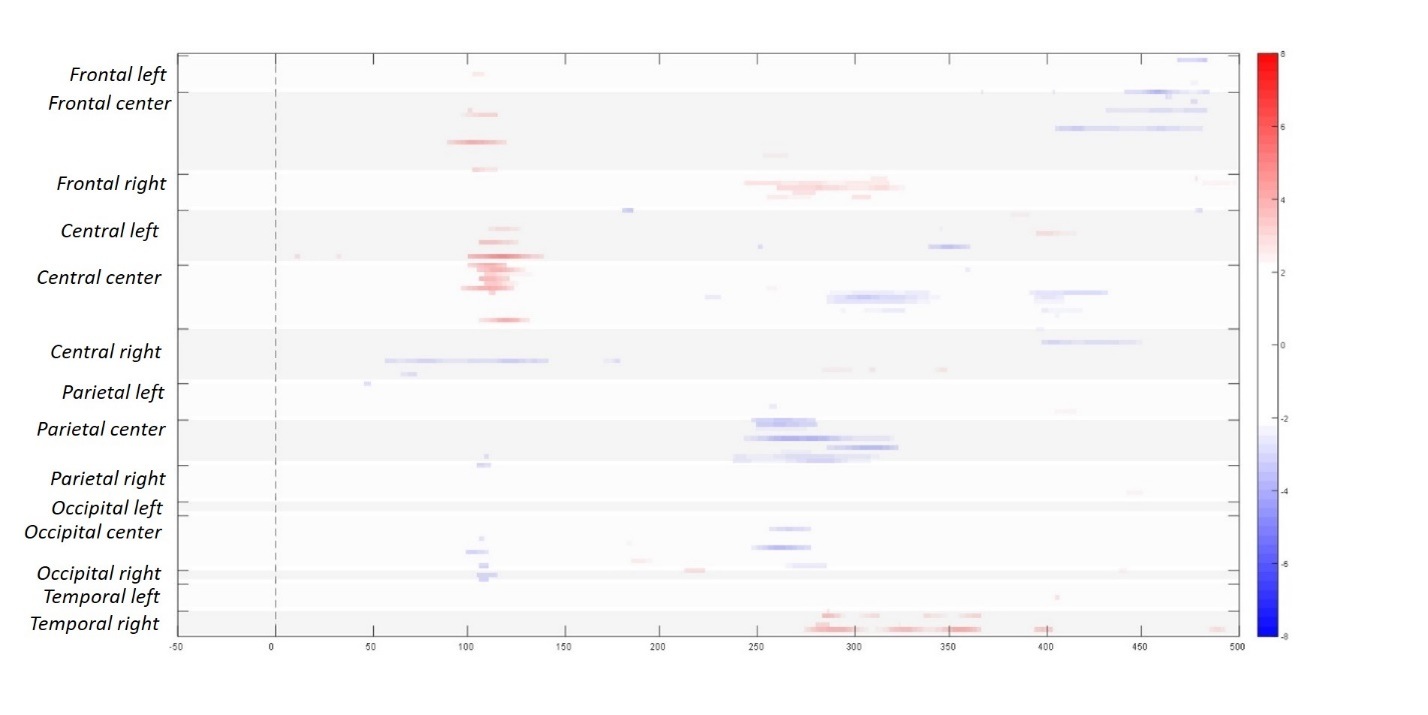
Supplemental Figure S6: T-values associated to the multiple t-tests analysis.

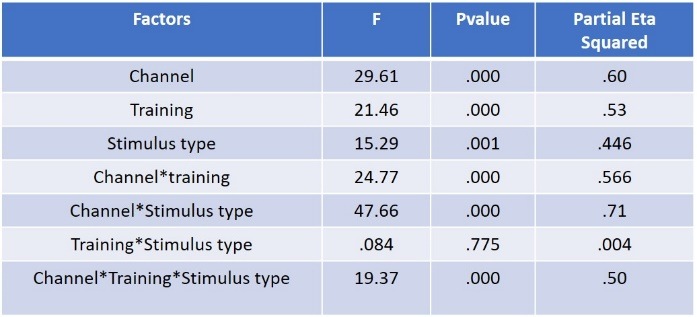

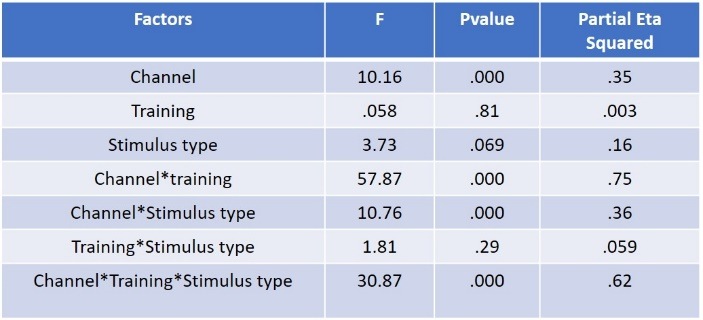

Table 4: Left panel: stage 2, matches Vs. mismatches (Cluster # 1). Comparison between the mean amplitude in the temporal window from 100 to 150 ms. Analysis performed on 20 participants. Right panel: stage 3, matches Vs. mismatches (Cluster # 2). Comparison between the mean amplitude in the temporal window from 250 to 300 ms. Analysis performed on 20 participants.

Based on this results, we defined these clusters as Regions of Interest (ROIs). Afterward, we run a power analysis, using the software G-Power, to establish the number of participants that needed to be tested to confirm the significance of these results. For the first temporal window (100-150 ms), considering an effect size equal to 1, α error probability equal to 0.05, one group and 40 measurments (10 x 2 x 2), the sample size was equal to 4. For the second temporal window (250-300 ms) given an effect size equal to 1.277, α error probability equal to 0.05, one group and 24 measurments (6 x 2 x 2), the sample size was equal to 3. We then collected data from 10 more participants and measured learning effects in our ROIs to determine the replicability of these effects.

After confirming the significant interactions between the factors, we reanalyzed data on all the 30 participants. Results are reported in table 6.

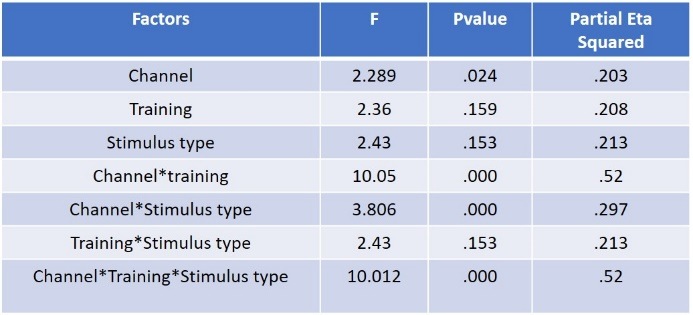

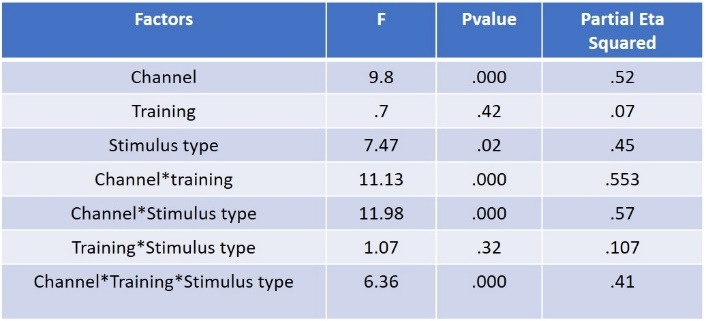

Table 5: Left panel: stage 2, matches Vs. mismatches (Cluster # 1). Comparison between the mean amplitude in the temporal window from 100 to 150 ms. Analysis performed on 10 participants. Right panel: stage 3, matches Vs. mismatches (Cluster # 2). Comparison between the mean amplitude in the temporal window from 250 to 300 ms. Analysis performed on 10 participants.

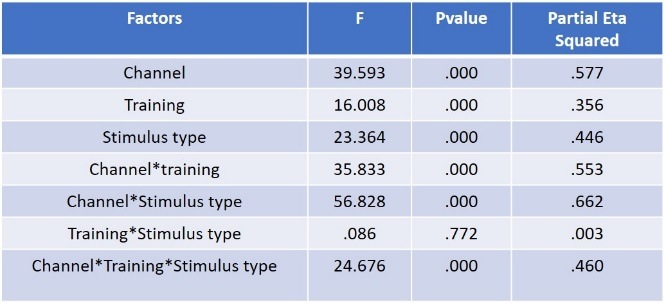

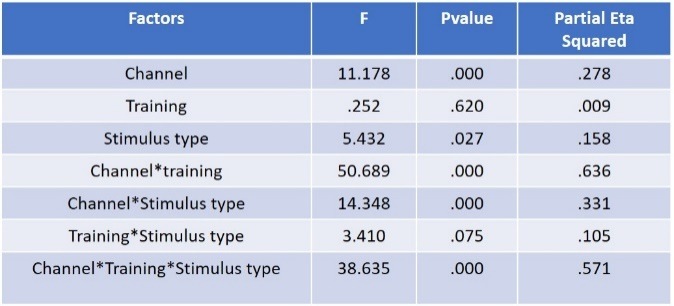

Table 6: Left panel: stage 2, matches Vs. mismatches (Cluster # 1). Comparison between the mean amplitude in the temporal window from 100 to 150 ms. Analysis performed on all the 30 participants. Right panel: stage 3, matches Vs. mismatches (Cluster # 2). Comparison between the mean amplitude in the temporal window from 250 to 300 ms. Analysis performed on all the 30 participants.

### **General multisensory effects**

Multisensory effects were measured as the difference between brain responses evoked by audiovisual stimuli and responses evoked by unisensory stimuli. Data were pooled across the two audiovisual conditions (matches/mismatches) and were then subtracted to the sum of the unisensory visual and audio responses. We considered the following channels: Oz, O1, O2, Pz, P1,P2, Fz, F1 and F2 and the following temporal windows: 80-100 ms, 130-150 ms and 250-500 ms. We ran repeated measures ANOVAs (one for each temporal window) with factors: channels, training (before/after) and stimulus type (multisensory/unisensory sum). Results from these analyses are reported in tables 7-9.

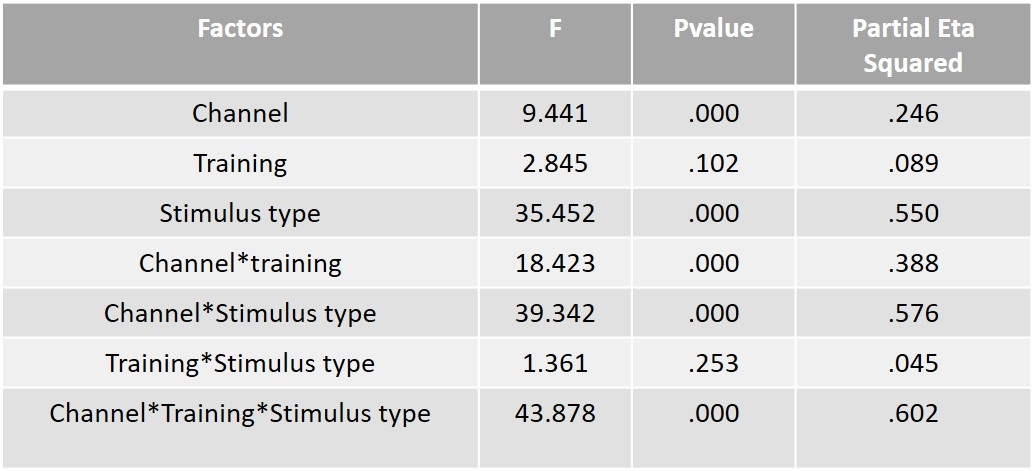

Table 7: Stage 1, multisensory Vs. unisensory sum. Comparison between the mean amplitude in the temporal window from 80 to 100 ms. Analysis performed on all the 30 participants.

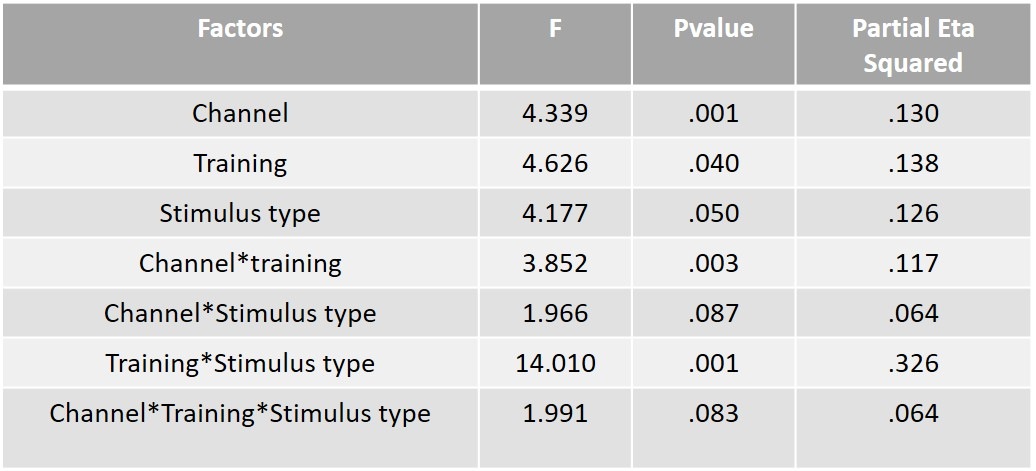

Table 8: Stage 2, multisensory Vs. unisensory sum. Comparison between the mean amplitude in the temporal window from 130 to 150 ms. Analysis performed on all the 30 participants.

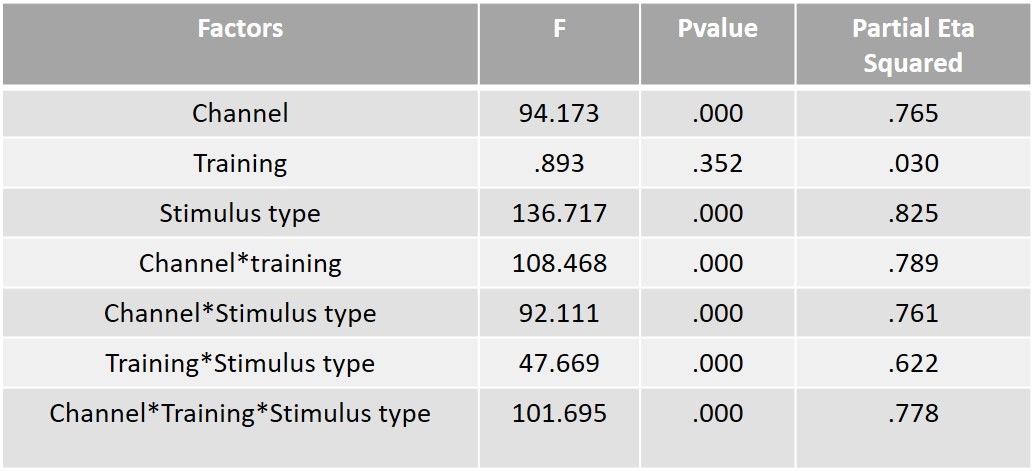

Table 9: Stage 3, multisensory Vs. unisensory sum. Comparison between the mean amplitude in the temporal window from 250 to 500 ms. Analysis performed on all the 30 participants.
